# Flexible coding schemes in dorsomedial prefrontal cortex underlie decision-making during delay discounting

**DOI:** 10.1101/2023.06.15.545101

**Authors:** Shelby M. White, Mitchell D. Morningstar, Emanuela De Falco, David N. Linsenbardt, Baofeng Ma, Macedonia A. Parks, Cristine L. Czachowski, Christopher C. Lapish

## Abstract

Determining how an agent decides between a small, immediate versus a larger, delayed reward has provided insight into the psychological and neural basis of decision-making. The tendency to excessively discount the value of delayed rewards is thought to reflect deficits in brain regions critical for impulse control such as the prefrontal cortex (PFC). This study tested the hypothesis that dorsomedial PFC (dmPFC) is critically involved in flexibly managing neural representations of strategies that limit impulsive choices. Optogenetic silencing of neurons in the rat dmPFC increased impulsive choices at an 8 sec, but not 4 sec, delay. Neural recordings from dmPFC ensembles revealed that, at the 8-sec delay, the encoding landscape transitions to reflect a deliberative-like process rather than the schema-like processes observed at the 4-sec delay. These findings show that changes in the encoding landscape reflect changes in task demands and that dmPFC is uniquely involved in decisions requiring deliberation.

## INTRODUCTION

Impulsivity is broadly defined as the predisposition to act prematurely without foresight^1, 2^ and is a primary feature of several psychiatric conditions; including substance use disorders (SUD) and schizophrenia^1–3^. Delay discounting (DD) is an established behavioral measure of cognitive impulsivity that measures the rate that a temporally delayed reward is devalued^4, 5^. Identifying the neural mechanisms that underlie DD has far reaching consequences, from inspiring novel treatment approaches for several psychiatric conditions to understanding decision-making.

Understanding the cognitive underpinnings of impulsivity requires investigating how decision-making changes when rewards are delayed in time. Models that span several levels of decision-making have conceptualized deliberation as a process whereby evidence is accumulated in a ramping-like manner and terminated at the choice^5–7^. As a complicated behavior is learned, strategies can be implemented that limit the need to deliberate between the available options de novo each time choice is made. Strategy use is effective at reducing impulsive choice^5, 8, 9^. Conversely, individuals rely on deliberation more when choices are difficult^10, 11^. Deliberative processes require effort^12^, are slower^12, 13^, and require the ability to weigh future choice options via prospection^7, 11^. Therefore, determining how strategies can be implemented and potentially disregarded during difficult decisions that require deliberation is critical to understand the neural basis of an impulsive choice.

A failure to use prospection and thereby make decisions without regard for future consequences is referred to as non-planning impulsivity^2, 4, 14^, which can be differentiated from impulsivity more generally^15^. Conversely, engaging in prospection reduces impulsive choices, which highlights this construct as a potential target for intervention^9, 16, 17^. Planning refers to policies enacted prior to deciding that will guide the action taken^18^, while prospection refers to mental time travel where one imagines the future in order to pre-experience an event^9, 15, 19^. Prospection has been formally implemented in DD models where an agent assigns subjective value to choice options via a cognitive search process to mentally imagine the options during deliberation^7^. Therefore, it is necessary to understand how neural networks in vivo implement strategies that may rely on planning and/or prospection in order to limit impulsive choices.

Several studies have examined the role of the rodent dorsal medial prefrontal cortex (dmPFC) in DD with mixed outcomes. Some studies find effects of manipulating this brain region on DD^20–24^, while others do not^25^. There is a strong rationale for examining dmPFC function in DD, as it is broadly implicated as critical for goal-directed decision-making. This includes monitoring the relationship between actions and outcomes^26–28^ and flexibly managing representations of strategies^29–31^. More generally, it is suggested that dmPFC is involved in integrating contextual information to facilitate the representations of goals or task rules^32^. During DD, neural activity in dmPFC has been implicated in signaling the need to change strategy^33–35^ as well as maintaining value representations^36, 37^. Moreover, dmPFC has been implicated in the initiation of deliberative sequences^11^, which require prospection in order to assign value to the delayed reward^7^. Finally, rats with several documented alterations in dmPFC function and neurochemistry lack behavioral correlates of strategy and are more impulsive than those that implement strategy^5, 38–43^. Taken together, these data motivate the need to identify the computations performed by dmPFC during a DD task to understand impulsivity.

To understand the computations that underlie the high-level, abstract features encoded in dmPFC, it is necessary to examine neural activity at the ensemble level^44–49^. With this approach, dmPFC ensembles have been shown to track the ongoing cognitive demands of a task^50–52^. Specifically, acquiring a new rule^46, 53^ or exposure to new context^47, 48^ is accompanied by “remapping” at the ensemble level^27, 48^. Further, recently developed dimensionality reduction approaches found flexible dynamics in PFC ensembles where linear dynamics transition to rotational dynamics, which was suggested to reflect decision commitment^6, 49^.

This study tested the hypothesis that dmPFC is critically involved in flexibly managing neural representations of strategy during DD. This hypothesis was tested in male rats performing a DD task, where neural activity was actuated via optogenetics and measured at the ensemble level via high density neural recordings.

## RESULTS

### Assessment of Planning Behavior and Strategy in Delay Discounting

Optogenetic inhibition and awake-behaving recordings were acquired in separate groups of rats and focused during a period prior to the choice (Figure 1A). Progression of an individual trial is described for optogenetic manipulations (Figure 1A, top) and awake-behaving recordings (Figure 1A, bottom). I-value reflects the number of pellets dispensed by the adjusting lever (Immediate) on a given trial (see methods). The impact of choice on i-value across an individual session is depicted in Figure 1B: Each immediate choice *decreases* and each delay choice *increases* the i-value on the subsequent trial by one. Forced trials have no impact on i-value (Figure 1B). Examples of strategy types (Immediate Exploit, Delay Exploit, titration) are also depicted in Figure 1B.

**Figure 1.**
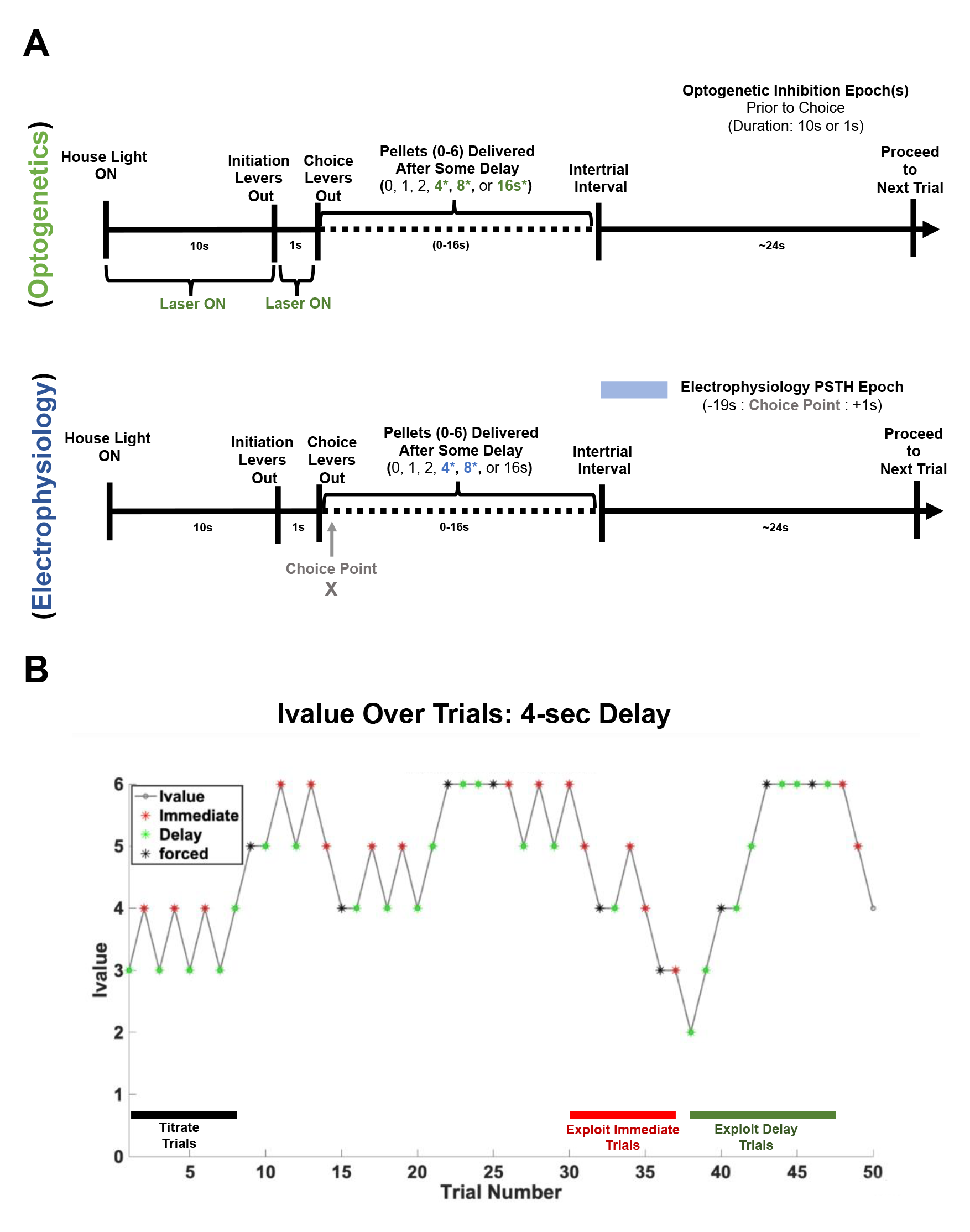
Description of optogenetics and electrophysiology experiments. **A)** Description of single-single trial during DD depicting optogenetic inhibition of dmPFC (**A**, top) and epochs selected for analysis in awake-behaving recordings from dmPFC (**A**, bottom). The green highlighted portion of the trial depicts the points during a single trial where laser was turned ON (**A,** top) while the blue highlighted region (**A**, bottom) depicts the portion of the trial analyzed for the electrophysiology experiment. **B)** Example single-session depicting how an animal makes choices during the 4-sec delay. Different strategies that emerge across trials are also depicted [Immediate-exploit (red), Delay-exploit (green), titration (black)]. Choice trials are depicted in red (immediate choices) and green (delay choices) while forced trials are shown in black. I-value refers to the number of pellets dispensed by the adjusting (Immediate) lever on a given trial.

### Delay choices are associated with correlates of planning behavior

Previous work from our lab suggests that planning is more prevalent on Delay choices than Immediate choices^5^. Planning would be expected to facilitate faster responses, therefore initiation and choice latencies were compared for Immediate and Delay choices in the absence of optogenetic stimulation (Figure 2A-D). Similar to our previous reports, Delay initiation latencies were shorter compared to Immediate (Wilcoxon Rank Sum test *Z*=6.21, *p*=4.98e-10; Figure 2A). In addition, Delay choice latencies were faster than Immediate choice latencies (Wilcoxon Rank Sum test *Z*=11.59, *p*=4.70e-31; Figure 2B). These data indicate that the animals plan Delay choices more than Immediate choices.

**Figure 2.**
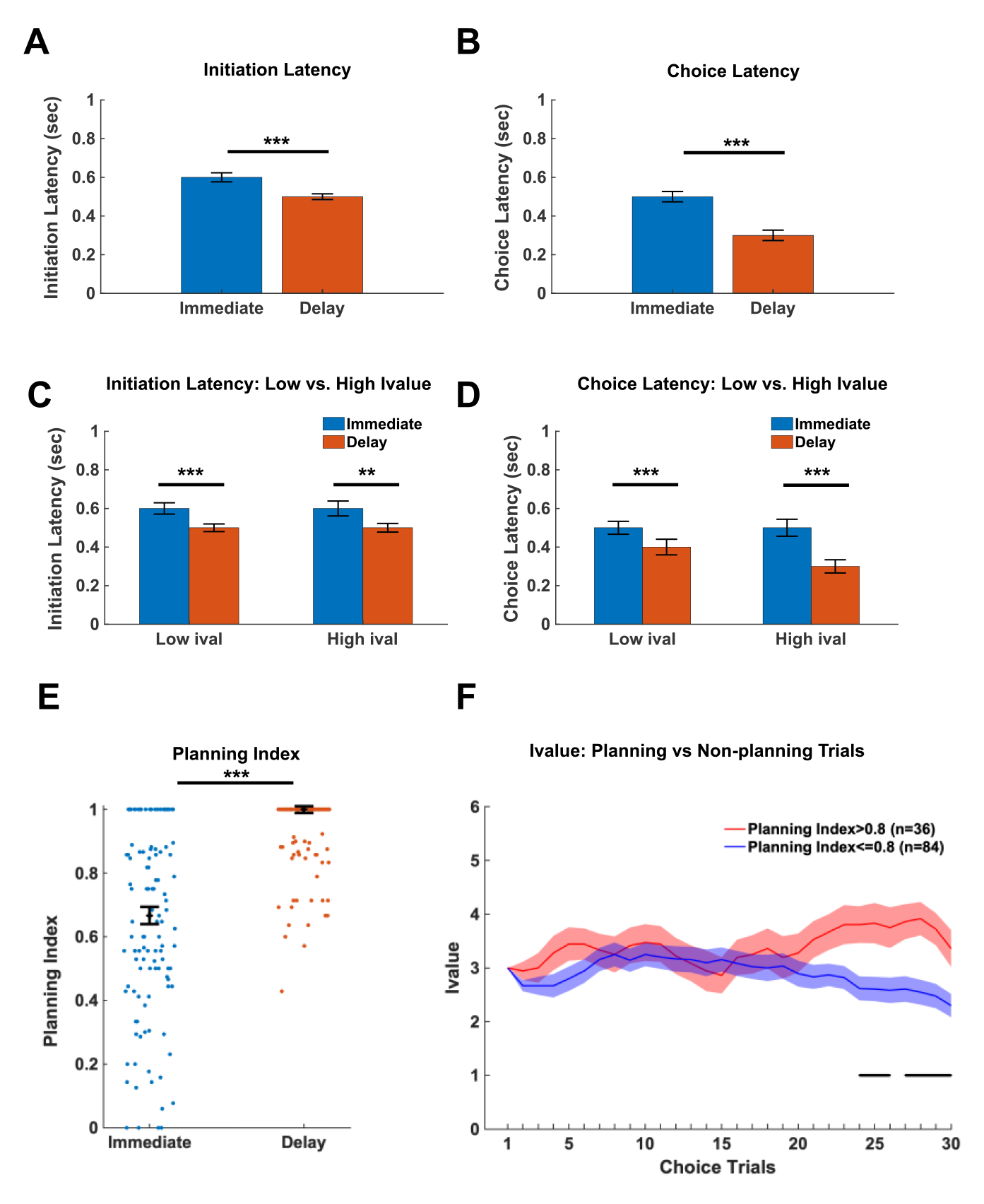
Choices are faster and more consistent when the Delayed option is selected. Assessment of latencies and planning index for optogenetic Laser OFF sessions (4, 8, 16-sec combined; n=8 animals). **A)** Delay initiation latencies (n=1864 choices) are shorter than Immediate (n=1736 choices) latencies (Wilcoxon Rank Sum: *Z=*6.22, *p* =4.98e-10). **B)** Delay choice latencies (n=1864 choices) are shorter than Immediate (n=1736 choices) latencies (Wilcoxon Rank Sum: *Z*=11.59, *p*=4.70e-31). **C)** Initiation latencies are longer for Immediate vs Delay choices for high i-value (>3) choices (Bonferroni-corrected Wilcoxon Rank Sum: *Z=*3.46, *p*=.002; n_Immediate_=899, n_Delay_=852 High i-value choices). Initiation latencies are also longer Immediate choices for low (<=3) i-value choices (Bonferroni-corrected Wilcoxon Rank Sum: *Z=*5.50, *p*=1.13e-7; n_Immediate_=837, n_Delay_=1012 Low i-value choices). **D)** Choice latencies are longer for Immediate vs Delay choices for high i-value (>3) choices (Bonferroni-corrected Wilcoxon Rank Sum: *Z=*7.91, *p*=7.70e- 15; n_Immediate_=899, n_Delay_=852 High i-value choices). Choice latencies are also longer Immediate choices for low (<=3) i-value choices (Bonferroni-corrected Wilcoxon Rank Sum: *Z=*8.38, *p*=1.56e-16; n_Immediate_=837, n_Delay_=1012 Low i-value choices). **E)** Planning index during Delay choices was higher than during Immediate choices (Wilcoxon signed-rank test: *Z*=-7.68, *p*=1.57e-14). Individual points show planning index for individual sessions (n=120 sessions) during Immediate or Delay choices. **F)** i-value (Mean +/- SEM) across choice trials was greater for high (red) than low (blue) planning index sessions (Wilcoxon Rank Sum test: Z=6.80, *p*=1.06e-11; n=36 high Planning index and n=84 low Planning index sessions). The solid line black line in **F** indicates points during the thirty choice trials where i-value differs for High vs Low Planning Index sessions for FDR-corrected individual Wilcoxon Rank Sum tests at each choice trial (n=30 choice trials). Median latencies (+/- SEM) plotted **A-E**. **P<.01, ***P<.001.

Given that the Immediate lever outcome fluctuates in response to previous choices, we investigated whether high (>3) or low (<=3) i-values influence latencies corresponding to Immediate or Delay initiations and choices. In all cases, Delay initiations (Figure 2C) and choices (Figure 2D) were faster than immediate (Bonferroni corrected Wilcoxon Rank Sum tests, *Z’s*>3.46, *p’s*<.01). No differences in initiation latencies were observed for high or low i-value conditions when collapsed across immediate and delay choices (Wilcoxon signed-rank test *Z*=.12, *p* =.90; Figure 2C). Similarly, when comparing choice latencies (Figure 2D), Delay choices were faster than Immediate choices for both low and high i-values (Bonferroni corrected Wilcoxon signed-rank tests *Z’s*>7.90, *p’s*<.001) and no differences were observed between high and low i-value choices when collapsed across Immediate or Delay (Bonferroni corrected Wilcoxon signed-rank test *Z*=-1.83, *p*=.07). Collectively, these results suggest that animals plan their responses on the Delay lever to a greater degree than the Immediate lever irrespective of i-value.

To further assess the hypothesis that Delay choices are planned to a greater degree than Immediate choices, we assessed patterns of responding between the initiation and choice lever for Delay and Immediate choices. For example, consistently using the same lever to both initiate and make a choice (i.e. initiation=Delay lever & choice=Delay lever for a given trial) suggests that the choice is being guided by a pre-existing plan. Alternatively, a minority of animals were observed to consistently initiate with the lever opposite to the choice lever (i.e. initiation on Delay lever and choice on Immediate). To quantify these two different planning types, a planning index was utilized. This value was multiplied by two to keep the range of values between zero (no indication of planning) and one (planning on every trial):

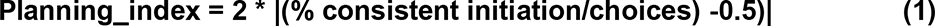

Providing further support for our hypothesis that animals tend to plan Delay choices to a greater degree than Immediate, a higher planning index was observed for Delay choices (Wilcoxon signed-rank test *Z*=-7.68, *p*=1.57e-14; Figure 2E).

The impact of planning on i-value was also assessed. I-value was higher for high (>.8) planning index compared to low (<=.8) planning index sessions (Wilcoxon Rank Sum test, *Z*=6.80, *p*=1.06e-11; Figure 2F), which was prominent at the end of the session (Figure 2F). Collectively, these results indicate that compared to Immediate choices, animals use a greater degree of planning for Delay choices which corresponds to decreased measures of impulsivity (higher i-value).

### Optogenetic inhibition of dmPFC increases impulsive choices at an 8-sec delay

Optogenetic inhibition of dmPFC was used to probe the impact of dmPFC activity on planning and impulsivity. It was hypothesized that inhibiting dmPFC would result in decreased planning, and consequentially, increase measures of impulsivity. Bilateral expression of ArchT in the dmPFC was present for all animals included in the analyses with majority expression in the dorsal mPFC with some ventral spread (Figure 3A). Indifference points for each condition (Laser ON vs Laser OFF) were calculated by averaging the last ten trials of each session for each delay (Figure 3B). A Mazur hyperbolic discounting function^54^ was fit to observations in the Laser ON and Laser OFF conditions (see supplement Figure 1). To assess differences between conditions, direct statistical comparisons of the two discounting curves an Extra-sum-of-squares F-test was used to determine whether one model accurately describes both conditions compared to the individual curves. One curve did not adequately fit both conditions (Extra-sum-of-squares F-test: F(1,46)= 10.46, *p* = .002; Figure 3B), indicating differences in *k*-values between conditions. Specifically, the Laser ON condition resulted in a steeper discounting curve compared to the Laser OFF condition (Figure 3B, left). Additionally, Laser ON vs OFF sessions were compared via Area Under the Curve (Figure 3B, right). Lower AUC for the Laser ON versus OFF was observed (paired samples t-test, *t*(7)=5.3, *p*<.01). Collectively these data indicate that inhibition of the dmPFC increases impulsive responding.

**Figure 3.**
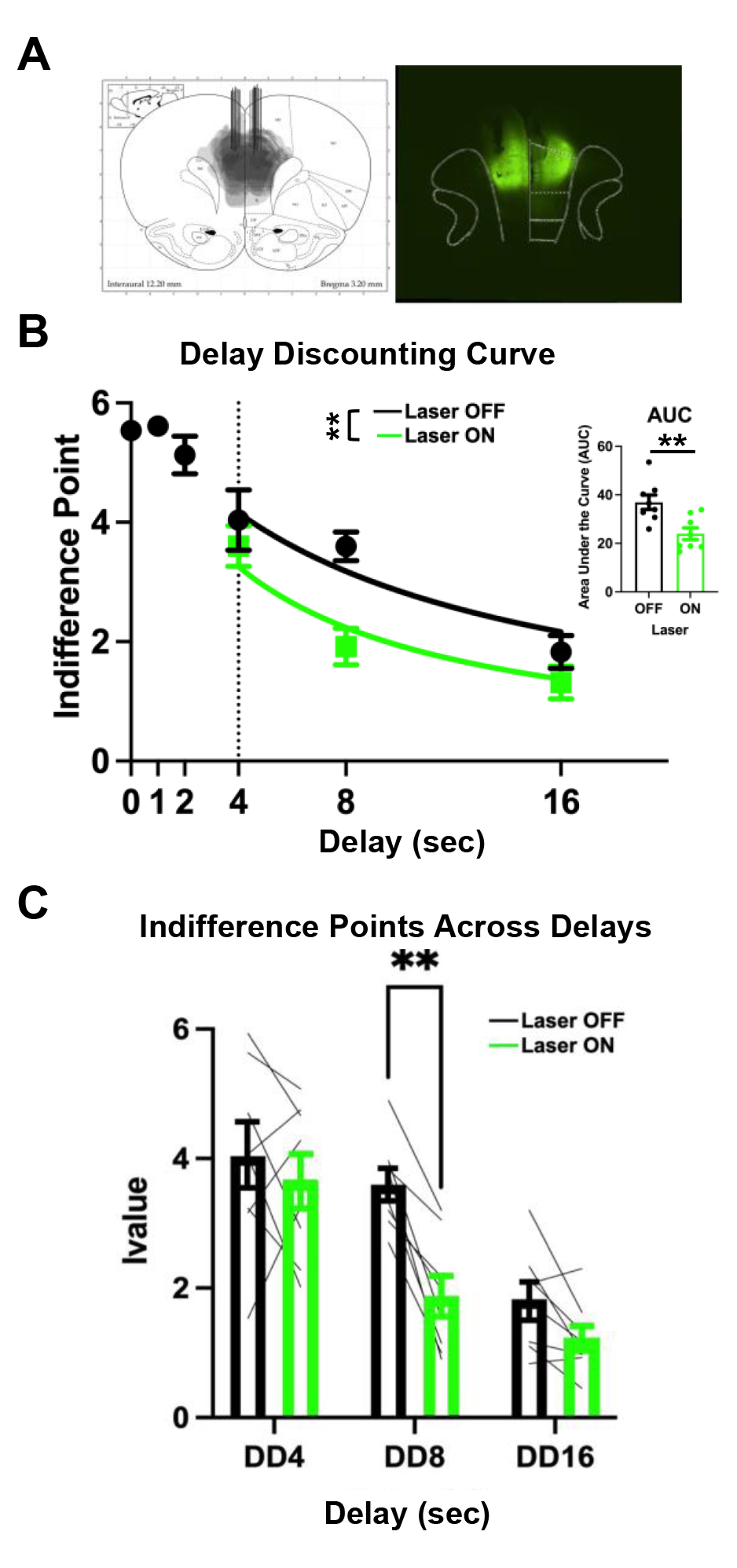
Optogenetic inhibition of dmPFC prior to choice increases impulsivity measures. **(A)** ArchT expression spread and optic fiber placements (left) for all animals (n=8). Representative image of viral spread and optic fiber placements (right). **(B)** Higher DD hyperbolic curve during Laser ON (green) than Laser OFF (black) sessions (4, 8, 16-sec) showing one curve did not adequately account for Laser ON and Laser OFF sessions. Area Under the Curve (AUC) corresponding the DD curve is greater for Laser OFF sessions than Laser ON sessions (paired samples t-test, *t*(7)=5.3, *p*<.01). **(C)** Indifference points decrease as delay increases (ANOVA: F(1.48,10.35)=27.53, p=.0001). Indifference points were decreased on Laser ON (green) sessions compared to Laser OFF (black) sessions (ANOVA: F(1,7)=27.53, p=.003). Specifically, at the 8-sec delay (Holm-Šídák test, *p*<.01) indifference points were decreased for Laser ON (green) compared to Laser OFF (black) sessions but not the 4-sec or 16-sec delays (Holm-Šídák tests, *p’s*>0.05). Optogenetic manipulation occurred at the 4-sec, 8-sec, and 16-sec delays **P< 0.01.

To further assess what drove the differences between the Laser ON and Laser OFF conditions, indifference points were compared at 4, 8, and 16s delays individually (Figure 3C). Greenhouse-Geisser corrected 2-way repeated measures ANOVA revealed a main effect of Delay (F(1.48,10.35)=27.53, p=.0001) and main effect of Laser ON/OFF condition (F(1,7)=27.53, p=.003); Figure 3C). Post hoc comparisons were then used to assess differences in impulsivity at each delay. While no differences were observed between Laser ON versus Laser OFF at the 4 or 16-sec delay (Holm-Šídák tests, *p’s*>0.05), at the 8-sec delay, the indifference point for the Laser OFF condition was larger than Laser ON condition (Holm-Šídák test, *p*<.01; Figure 3C).

These results indicate that the indifference points between Laser ON/OFF conditions during the 8-sec delay were the major factor in differences between impulsivity measures. The selective effect at the 8-sec delay may reflect the relatively equal probabilities of choosing either the delay or immediate lever (supplemental Figure 3).

To investigate the impact of optogenetic inhibition on planning index during the last ten choice trials of the session (indifference points), Laser ON vs Laser OFF conditions were compared across all three experimental delays (4, 8, 16-sec). Contrary to our hypothesis, optogenetic inhibition had no impact on planning index across each delay (Figure 4A). This was true regardless of choice type (Immediate or Delay; data not shown).

**Figure 4.**
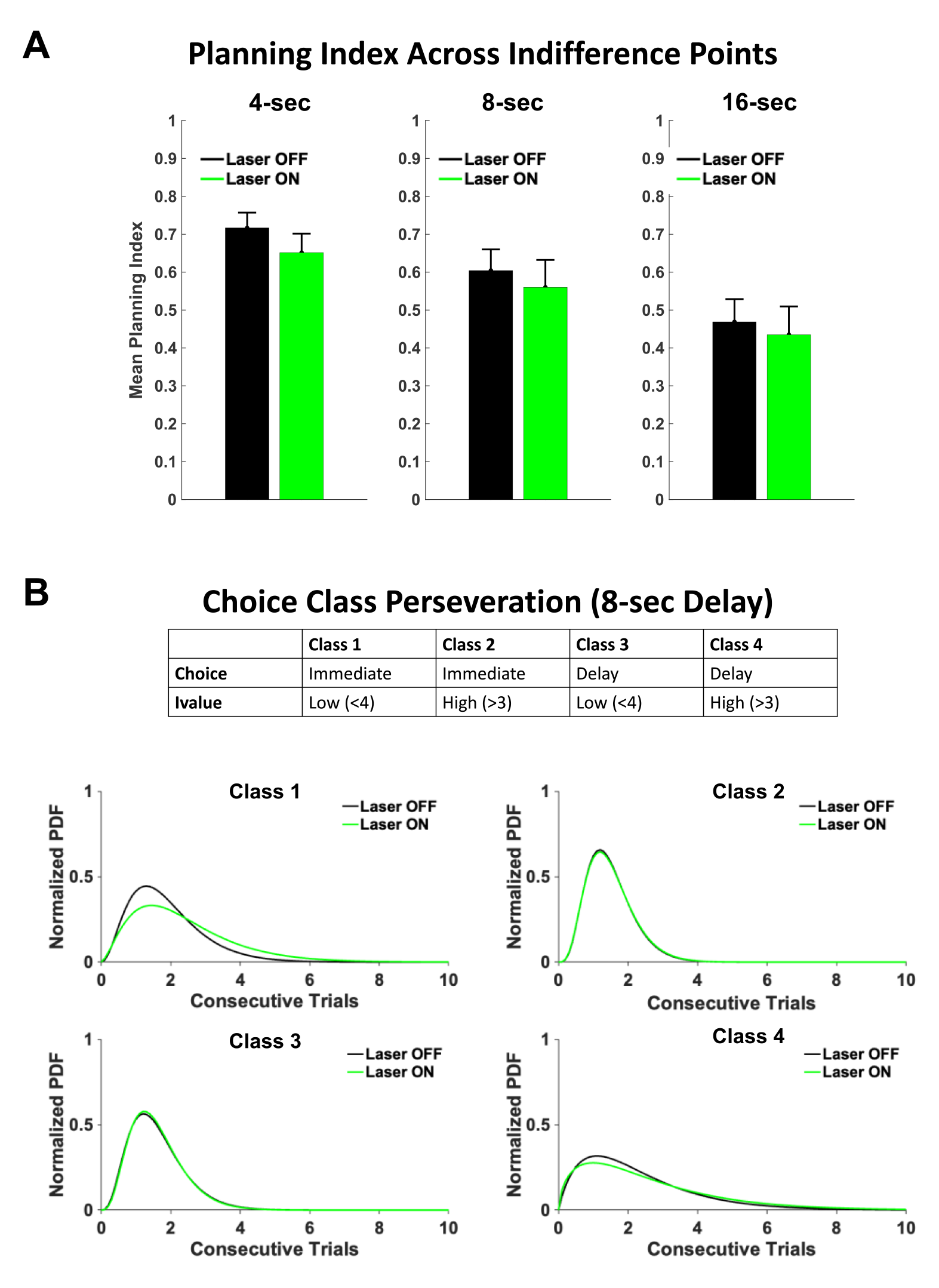
Optogenetic inhibition of dmPFC disrupts strategy change that occurs over trials rather than planning of individual choices. **(A)** Optogenetic inhibition of dmPFC (n=8 animals) did not lower planning index for the 4-sec (*t*(52)=-1.01, *p*=.31), 8-sec (*t*(50)=-0.49, *p*=.63), or 16-sec (*t*(53)=-0.36, *p*=.72) delay. **(B)** Decisions separated into choice classes (B, top) shows optogenetic inhibition increases the number of class 1 choices at the 8-sec delay (Kolmogorov-Smirnov: *p*=0.022).

The hypothesis was then tested that dmPFC may contribute to strategies that impact choice behavior on longer time scales (i.e., across trials). To examine choice sequences, trials were split into 4 different classes (Figure 4B, top) based on i-value and choice type. This assessed the ability of the animal to deviate from poor choice strategies, such as exploiting the Immediate lever when the i-value is low. Class 1 and Class 4 decisions were considered the least optimal and we hypothesized that optogenetic inhibition would disrupt the ability to switch from exploit strategies in these instances. For both the Laser ON and Laser OFF conditions we determined the distribution of consecutive choices within each class. A gamma-distribution was fit to both the Laser ON and OFF data. Laser ON conditions lengthened consecutive Class 1 decisions in the 8-sec delay (Kolmogorov-Smirnov, p=.022, k=.32; Figure 4B). This indicated that optogenetic inhibition disrupted dmPFC signals required to shift away from poor decisions. No effects of inhibition were seen in the 4-sec delay. Collectively these results suggest that, at the 8-sec delay, optogenetic inhibition of dmPFC increases impulsivity by impacting the animals’ ability to deviate from exploiting the Immediate lever when the value is low.

### dmPFC networks shift from schema encoding at 4-sec to deliberative encoding at 8-sec

To determine why optogenetic inhibition of dmPFC increased impulsivity at the 8-sec (but not 4 sec) delay, male Wistar rats were unilaterally implanted with 64-channel silicone probes in dmPFC (see methods; Figure 5A) and recordings during the 4- and 8-sec delay were analyzed. To understand the underlying differences in neural activity patterns for Immediate and Delay choices generally, the mean firing rates for Immediate and Delay choices were aligned to 15-sec prior to the choice and 15-sec after and were plotted for the 4-sec and 8-sec delay. Mean firing rates of choice options exhibited ramping activity prior to the choice point for Immediate and Delay choices and were shifted for 8-sec choices compared to 4-sec choices (Figure 5B, C).

**Figure 5.**
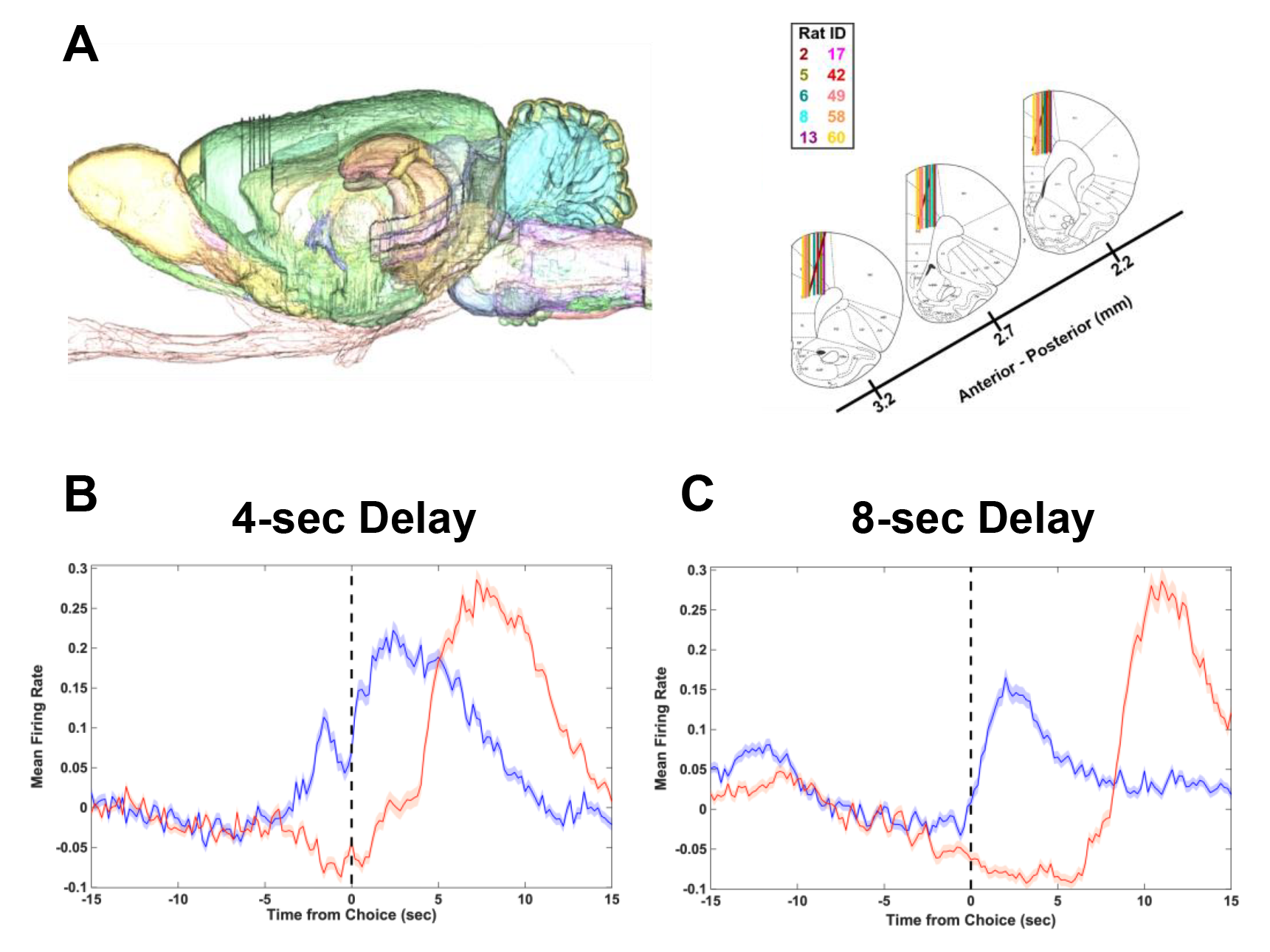
Neural activity of dmPFC neural populations during Immediate and Delay choices during the 4 sec and 8 sec delay. (A) Electrophysiology placements of silicone probes. Representative image of sagittal slice with probe placement in the right dmPFC (left) and for all animals (right). **(B-C)** Grand average mean firing rate for the 4s **(A)** (n=2120 neurons) and 8s **(B)** delay (n=2078 neurons) separated by Immediate (blue) and Delay (red) choices aligned to the time that the animal presses the choice lever (dashed line at time=0).

Given that optogenetic inhibition occurred prior to the choice point, the remainder of the analyses were conducted on spike trains 19-sec prior to and 1-sec after the choice was made on a given trial (Figure 1A). Additionally, to better understand neural activity related to strategy shifts, the remainder of the analyses focused on trials that occur specifically during exploitation of the Immediate or Delay lever. Immediate and Delay Exploit strategies (Figure 1A) were defined as three or more consecutive choices on the same lever (Immediate or Delay), given that animals would have to continue pressing on the third trial despite being exposed to a forced trial. Therefore, neural activity was assessed between the 3^rd^ and 4^th^ choice trial in the sequence. The 4^th^ choice could either be to continue to exploit (Fail-to-Change strategy) or to shift to the opposite lever (Change strategy). The four types of strategy shifts analyzed were a ‘IM Change’, ‘IM Fail-to-Change’, ‘DEL Change’, and ‘DEL Fail-Change’ strategy (see Figure 6A, top). The four strategy types enabled the assessment of whether neural signatures of continuing to exploit one choice (IM/DEL Fail-Change) differed from those of choosing to abandon an exploit strategy on the fourth trial (IM/DEL Change). To test this, PCA was performed across each strategy type on neurons for the 4-sec (n=581 neurons) and 8-sec (n=1166 neurons) delays to obtain the neural trajectories of the 3^rd^ and 4^th^ trial (see Materials and Methods).

**Figure 6.**
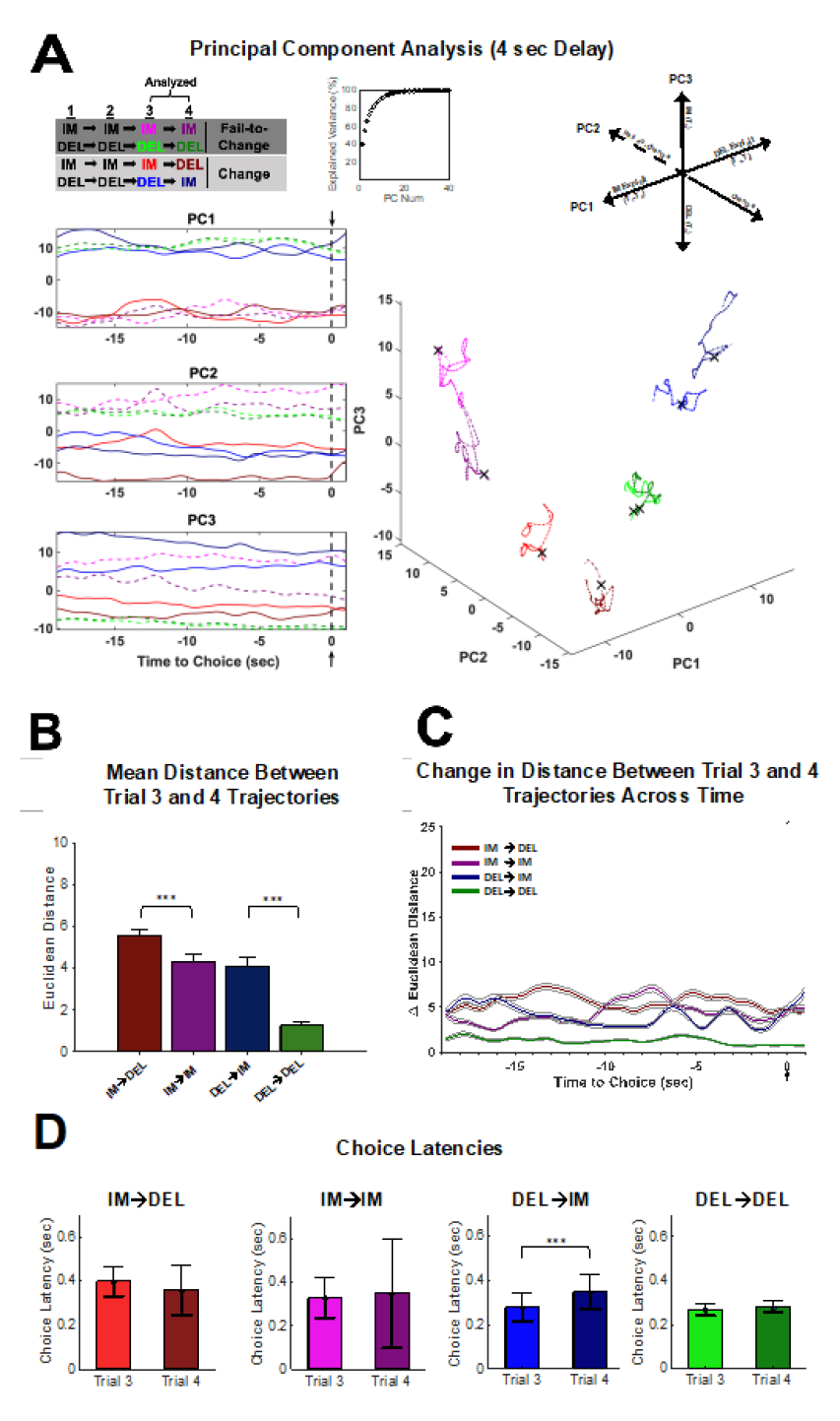
PCA analysis for strategy transitions during the 4-sec delay reveal schema-guided decision-making. **(A)** Population activity (n=581 neurons across n=11 sessions) during the 3^rd^ and 4^th^ trial of a four-trial sequence for all four strategy shifts (IM-Change, IM-Fail-to-Change, DEL-Change, DEL-Fail-to-Change) was analyzed using PCA. Trials 1-3 (T_1-3_) were either IM or DEL exploit trials and Trial 4 (T_4_) was either the ‘Change’ or ‘Fail-to-Change’ trial (see key: A, top left). Top 3 PCs showing trajectories for trial 3 and trial 4 of the four strategy shifts analyzed plotted for each individual top PC (A, left). Dashed lines in the individual PC plots depict Fail-to-Change conditions while solid lines indicate Change conditions (A, left). Cumulative explained variance for the top 40 PCs (A, top middle). Evolution of trajectories for trials 3 and 4 for each strategy condition in 3D PC space for the top 3 PCs, with each dimension corresponding to PC1, PC2, or PC3 (A, bottom right). The **X** denotes the choice point in the trajectory (A, bottom right). Summary of how encoding of task variables relevant to strategy change occurs in 3D PC space (from top 3 PCs), revealing coding schema (A, top right). **(B)** Euclidean distance (mean +/- SEM) between the third and fourth trial trajectory of each strategy shift. The average distance between trials 3 and 4 trajectories did not significantly differ between IM-Fail-to-Change and DEL-Change conditions (Tukey Kramer: *p*=.84) but all other comparisons of strategy shifts were significantly different from one another (Tukey Kramer: *p*<.001). **(C)** Change in Euclidean distance (mean +/- SEM) between trials 3 and 4 across time showing how the distance between trial 3 and 4 differs leading up to the choice point (dashed gray line) for each of the four strategy transitions. PCA conducted on individual sessions for distance measures (B-C; n=11 sessions, see Material and Methods). **(D)** Choice Latencies compared for trials 3 and 4 for each of the four strategy shifts. Choice latencies increase on trial 4 compared to trial 3 only when the animal shifts away from the exploiting the Delay lever (Wilcoxon Signed Rank test: *Z*=-3.74, *p*=.0002). No differences in choice latencies were observed between trial 3 and 4 for any other strategy shift (Wilcoxon signed-rank test: *Z*’s<1.72*, p*’s>.05). ***P<.001.

The top three PCs and the trajectory of each strategy type are plotted for the 4-sec (Figure 6A) and the 8-sec delays (Figure 7A). For the 4-sec delay, each of the first 3 PC’s clearly separated features of the task (Figure 6A); where PC1 reflected neural activity patterns related to the exploit sequence in which the animal was engaged (IM-IM-IM vs DEL-DEL-DEL), PC2 separated if the animal would continue to exploit or not (Change vs Fail-to-Change), and PC3 separated what choice the animal made on the 4^th^ trial (IM vs DEL). At the 8-sec delay, PC1 still reflected the exploit sequence (Figure 7A). However, PC2 no longer reflected exploit or not, and PC3 reflected DEL-change or DEL-Fail-to-Change sequences (Figure 7A). These data suggest that the encoding schemes in dmPFC differ across the delays of the DD task.

**Figure 7.**
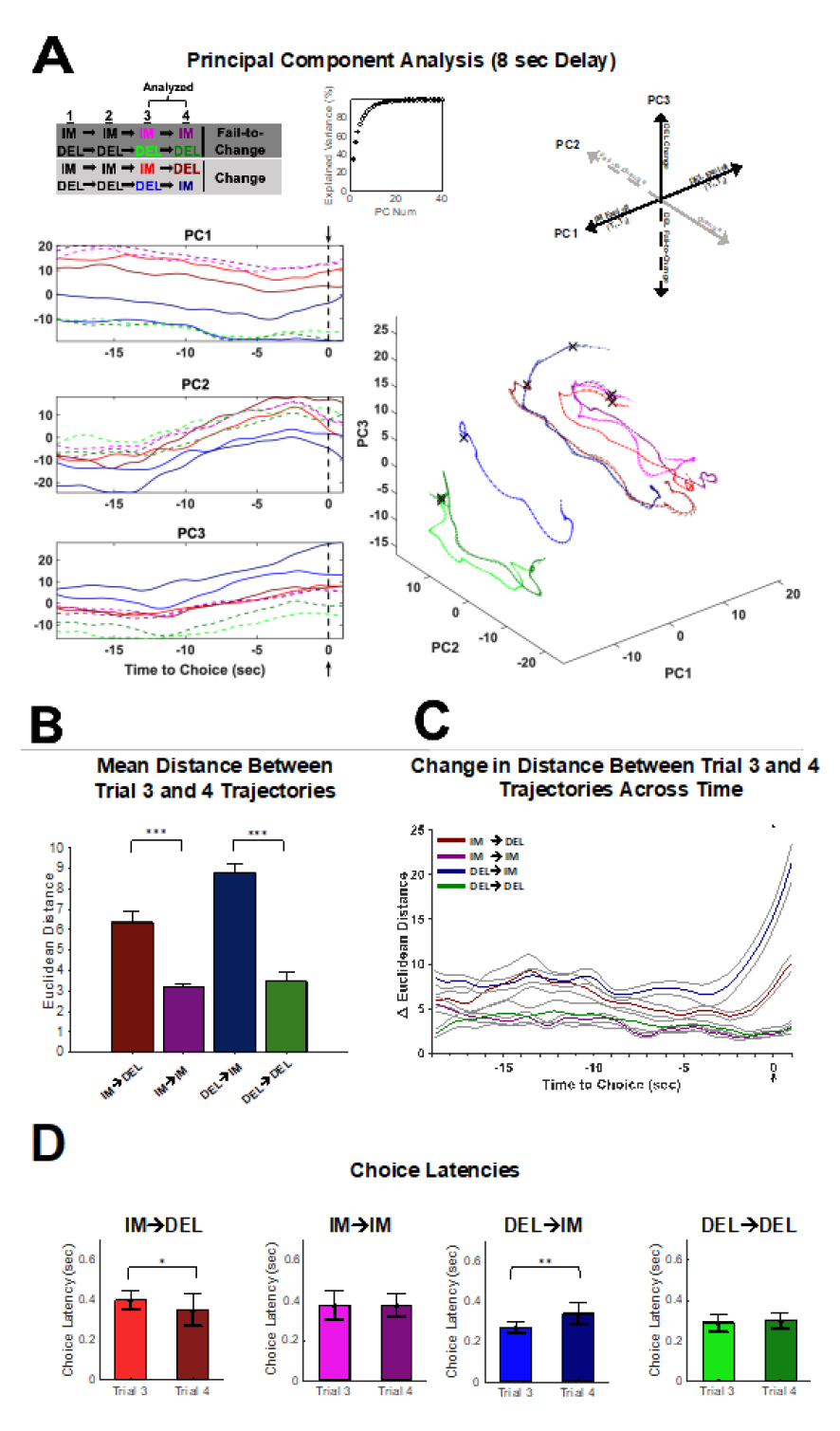
PCA analysis for strategy transitions during the 8-sec delay reveal deliberation-guided decision-making. **(A)** Population activity (n=1166 neurons across n=13 sessions) during the 3^rd^ and 4^th^ trial of a four-trial sequence for all four strategy shifts (IM-Change, IM-Fail-to-Change, DEL-Change, DEL-Fail-to-Change) was analyzed using PCA. Trials 1-3 (T_1-3_) were either IM or DEL exploit trials and Trial 4 (T_4_) was either the ‘Change’ or ‘Fail-to-Change’ trial (see key: A, top left). Top 3 PCs showing trajectories for trial 3 and trial 4 of the four strategy shifts analyzed plotted for each individual top PC (A, left). Dashed lines in the individual PC plots depict Fail-to-Change conditions while solid lines indicate Change conditions (A, left). Cumulative explained variance for the top 40 PCs (A, top middle). Evolution of how trajectories for trials 3 and 4 for each strategy condition move in 3D PC space for the top 3 PCs, with each dimension corresponding to PC1, PC2, or PC3 (A, bottom right). The **X** denotes the choice point in the trajectory (A, bottom right). Summary of how encoding of task variables relevant to strategy change occurs in 3D PC space (from top 3 PCs), revealing coding schema (A, top right). **(B)** Euclidean distance (mean +/- SEM) between the third and fourth trial trajectory of each strategy shift. The average distance between trials 3 and 4 trajectories did not significantly differ between Fail-to-Change conditions (Tukey Kramer: *p*=.55) but all other comparisons of strategy shifts were significantly different from one another (Tukey Kramer: *p*<.001). **(C)** Change in Euclidean distance (mean +/- SEM) between trials 3 and 4 across time showing how the distance between trial 3 and 4 differs leading up to the choice point (dashed gray line) for each of the four strategy transitions. PCA conducted on individual sessions for distance measures (B-C; n=13 sessions, see Material and Methods). **(D)** Choice Latencies compared for trials 3 and 4 for each of the four strategy shifts. Choice latencies significantly differed between trials 3 and 4 for the DEL-Change (Wilcoxon signed-rank test: *Z*=-3.16, *p* =.002) and IM-Change (Wilcoxon signed-rank test: *Z*=2.37, *p*=.02) strategy conditions. No differences in choice latencies were observed between trial 3 and 4 for the Fail-to-Change strategy conditions (Wilcoxon signed-rank test: *Z*’s<.34, *p*’s>.73). *P<.05, **P<.01, ***P<.001.

To quantify how encoding differs when an animal continues to exploit vs deviating from an exploit strategy, the mean Euclidean distance between each point in the 3^rd^ trial’s trajectory was compared to each point in the 4^th^ trial’s trajectory for each individual session (Figure 6B, Figure 7B). For both the 4-sec and the 8-sec delay, distances for each Change trial type (e.g. IM→DEL, DEL→IM) were larger than their respective Fail-to-Change types (e.g. IM→IM, DEL→DEL) (Kruskal-Wallis(3,403) *χ*2=270.39, *p*<.001, Figure 6B; Kruskal-Wallis(3,403) *χ*2=304.78 *p*<.001, Figure 7B). This indicates that the neural activity patterns across Change trials 3 and 4 differed more so than Fail-to-Change trials, which may reflect neural activity patterns required to update choice strategy. However, this was common across the 4 and 8-sec delays, therefore difficult to reconcile with the selectivity of the optogenetic inhibition at the 8-sec delay.

The change in distance leading up to the choice between the 3^rd^ and 4^th^ trials was analyzed (Figure 6C and 7C). An interaction between strategy conditions and time was observed in both the 4-sec (Repeated Measures ANOVA F(100,300)=2.956, *p*<.001; Figure 6C) and 8-sec (Repeated Measures ANOVA, F(100,300)=7.130, *p*<.001; Figure 7C) delays. Examination of the temporal patterns of activity captured in the PCs indicated that, at the 4-sec delay, they were relatively stable throughout the trial (Figure 6A). This was quantified in the trajectory differences between the 3^rd^ and 4^th^ trials (Figure 6C) at the 4-sec delay, which did not seem to systematically vary leading up to the choice. The stability of the PCs in time indicates that neural activity encoded something akin to a state variable rather than a dynamic process that might be expected of deliberation preceding a choice. Encoding of a state-like variable was also consistent with the information content that was detected in each PC (Figure 6A).

At the 8-sec delay, neural activity patterns exhibited ramping-like activity leading up to the choice which was most prevalent in PC2 but visually appreciable in each PC (Figure 7A). Ramping-like activity was quantified in the trajectory differences where Change sequences (e.g., IM→DEL, DEL→IM) exhibited ramping-like activity leading up to the choice, while Fail-to-Change trials were static (Figure 7C). These data indicate that, on Change trials, dmPFC neural ensembles exhibit a temporal profile that is consistent with a process like deliberation, where ensembles progressively evolve toward the choice point. This is consistent with the need to deliberate at the 8-sec delay as the probability of choosing either option is roughly equivalent (supplemental Figure 3).

If deliberation was more prevalent at the 8 sec than 4 sec delay, then changes in response latencies should be observed between the 3^rd^ and 4^th^ trials. At the 4-sec delay, animals increased choice latencies between the 3^rd^ and 4^th^ trial for DEL-Change sequences (DEL→IM; Wilcoxon signed-rank test: *Z*=-3.74, *p*=.0002; Figure 6D) but not for Fail-to-Change or IM-Change sequences (Wilcoxon signed-rank tests: *Z*’s<1.72, *p*’s>.05). This indicates that latencies increase only when switching away from the preferred (delay) lever (supplementary Figure 3) at this delay. In contrast, at the 8-sec delay, choice latencies differed between the 3^rd^ and 4^th^ trial for both the IM-Change (Wilcoxon signed-rank test: *Z*=2.37, *p*=.02) and the DEL-Change (Wilcoxon signed-rank test: *Z*=-3.16, *p*=.002) sequences but not for either Fail-to-Change sequences (Wilcoxon signed-rank tests: *Z*’s<.34, *p*’s>.73; Figure 7D). Responses were faster when going from IM→DEL and slower when going from DEL→IM and therefore consistent with the animals either switching to or away from their preferred option, which is consistent with deliberating an easy or difficult choice, respectively. In addition, this suggests that the effect of the optogenetic inhibition at the 8-sec delay interferes with the deliberative process that emerges across dmPFC ensembles.

Theories of DD propose that to evaluate the delay option the magnitude of the reward must be considered in the future context that it will be received. This is hypothesized to rely on a process of “cognitive search” where, as rewards are more temporally delayed, neural trajectories required to arrive at a representation of the value of the future reward are longer. We tested this hypothesis in our data as, if the reward was devalued after 8-sec delay because it was longer, the neural trajectories should cover a larger distance in state-space. This hypothesis was supported, as trajectories at the 8-sec delay were longer than those at the 4-sec delay (Mann-Whitney U test: U=1955, *p*<.001; c.f. Figure 6A, 7A) therefore supporting the observation that deliberative coding scheme is required for the 8-sec but not 4-sec delay.

## DISCUSSION

The main finding of this study is that when a clear choice preference exists (4-sec delay) the neural encoding landscape reflects a schema-driven decision process that changes to a deliberative-driven decision process when a choice preference no longer exists (8-sec delay). This was supported by changes in reaction time when animals update strategies concerning their preferred choice (4-sec delay) or both choice options (8-sec delay). Optogenetic inhibition of dmPFC increased impulsivity measures and increased the number of consecutive low-value choices at the 8-sec delay, suggesting that disrupting the deliberative encoding landscape increases measures of impulsivity by preventing strategy change signals. Collectively these data support the conclusion that the encoding landscape shifts based on the cognitive demands of the task and that impulsive choices are the result of a failure to engage a deliberative process to guide decision-making.

The differences in dmPFC ensemble activity that guide decision-making surrounding strategy change between delays were uncovered by examining PC spaces. When a clear preference for the Delay lever exists (4-sec delay; supplementary Figure 2), neural activity across dmPFC PC spaces were relatively static leading up to the choice. This indicates the existence of a latent signal in dmPFC ensembles that encoded features of the task in a static manner and that existed prior to optogenetic inactivation, which occurred 10 seconds prior to the choice. This may explain the lack of effect of the optogenetic inhibition at the 4-sec delay – any information that dmPFC might contribute to the decision at this delay preceded the inactivation. In addition, a schema encoding strategy may provide a cognitively economical way to choose that limits the need for cognitive resources (e.g., effort, prospection) and facilitates a more procedural way to arrive at a good decision.

In contrast, at the 8-sec delay, PC spaces were more dynamic surrounding strategy changes prior to the choice (i.e., during the window of time that dmPFC was inhibited via optogenetics). These data are consistent with the view that, during DD, dmPFC plays a critical role in signaling the need to change strategy^34^. Our data support and extend this view by indicating that that dmPFC may be uniquely involved when decisions require deliberation.

A number of studies have indicated that dmPFC is critically involved in decision-making when decisions are difficult^33, 55^. More deliberation is required for difficult decisions^10, 11^ and less deliberation occurs for difficult decisions when dmPFC is inhibited^11^. One explanation for animals failing to change strategy is that optogenetic inhibition of dmPFC disrupted deliberative processes that were more prevalent during the 8-sec delay. Decisions between the Immediate and Delay choice are most difficult at the 8-sec delay as, at this delay, the number of Immediate and Delay choices are roughly equal^5^ (supplementary Figure 2).

The behavior of the trajectories in the PC spaces provide insight into how dmPFC might implement the computations that control decision-making at each delay. While the differences across each delay were robust, a limit to making inferences about these spaces is the shortcoming of PCA. While useful for dimensionality reduction, PCA may not be sufficient to capture dynamics that occur in a high dimensional space given the linear nature of the algorithm^56^. Nonetheless, several important inferences have been made using PCA about latent dynamics that are supported by analysis tools better equipped to describes dynamics in a high-dimensional space.

The differences in the PC spaces across delays suggest that each delay exhibits a different degree of stability. The neural trajectories for each of the trial types at the 4-sec delay were restricted to a discrete region of state space that were generally well-separated from other trial types, suggesting the existence of several meta-stable states (e.g., multistability). Therefore, this suggests that multistability facilitates schema encoding strategies where decision-making is more procedural.

Prior work from our group indicates that neural trajectories in dmPFC track task variables by itinerating through meta-stable states during a foraging-based decision-making task^57, 58^ which is reminiscent of the encoding scheme observed at the 4-sec delay. This type of encoding scheme seemed, initially, to be observable at the 8-sec delay. However, at ∼12 seconds prior to the choice, PC spaces begin to evolve where PC’s 2 and 3 each begin to move upward (Figure 7A). The change in the way neural trajectories move through state space suggests that a bifurcation may occur and alter the dynamics that control dmPFC networks around this time. After this time, neural trajectories seem to take linear paths that give way to rotational paths near the choice. This type of linear to rotational dynamics is reminiscent of that observed in prefrontal networks of the nonhuman primate, which has been suggested to correspond to decision commitment^6, 49^.

Our data support that discounting emerges as a result of a cognitive search process^5, 7^. This theory proposes that choice options are evaluated through a process of prospection where options that are ‘easier to find’ are more likely to be chosen and options that are ‘more difficult to find’ are less likely to be chosen. In support of this theory, our data show that the distance between the start of a trajectory and the choice point is shorter (i.e., easier to find) for the 4-sec delay compared to the 8-sec delay regardless of the trial type (Figure 6A, Figure 7A).

The prospective aspect of cognitive search has been conceptualized in the framework of attractor dynamics^7^. Specifically, choice history impacts current choice behavior by imposing a bias that results in decision representations settling into basins of attraction that result in the same choice as what was chosen on the prior trial^59^, possibly because it was ‘easier to find’^7^. Therefore, disrupting dmPFC activity during deliberative processes, where the Immediate and Delay choice options must be weighed, has the potential to negatively impact the ability to ‘find’ the Delay option during prospection. The inability to find the Delay choice option would therefore be hypothesized to result in continued choice of the Immediate option, as was seen with optogenetic inhibition of the dmPFC. This rationale is supported by a recent study that shows that optogenetic *activation* of dmPFC prior to choice during a spatial DD task caused animals to take on a more deliberative rather than procedural strategy^24^, whereas our data show that optogenetic *inhibition* disrupts ability to use deliberative decision-making to implement necessary strategy changes.

Collectively, these observations form the basis of hypotheses that can be directly tested with modern techniques to reconstruct latent dynamics from neural recordings^46, 50, 52, 60^. Specifically, if a bifurcation exists and options are encoded via attractor dynamics, characterizing this will provide insight into how deliberation is implemented in dmPFC networks and how breakdowns in computation responsible for deliberation negatively impact decisions. Identifying methods to repair these computations, such as improving the ability to ‘find’ delay rewards during deliberation could be a powerful approach to reduce impulsivity and thereby improve treatment outcomes in several psychiatric disorders.

## METHODS

Methods and Supplemental materials and any associated references are available in the **online version of the paper**.

## ACKNOWLEGMENTS

We would like to thank Amanda Callahan for their assistance in conducting the optogenetic experiment. The work was supported by NIH grants AA029409, P60-AA007611, and T32AA007462.

## AUTHOR CONTRIBUTIONS

S.W. conducted and designed the optogenetic experiment. D.N.L, M.D.M., C.C.L, and C.L.C designed the *in vivo* recording study. S.M.W., M.D.M., B.M. and D.N.L. conducted the *in vivo* recordings. S.M.W., M.D.M. and E.D. conduced the data analyses on *in vivo* recordings. S.M.W., M.D.M., and C.C.L. wrote the paper.

## Supplementary Material

## METHODS

### Animals

Male Wistar rats were purchased from Envigo (Indianapolis, IN) to complete Experiment 1 (optogenetic inhibition of dmPFC; n=8) and Experiment 2 (awake-behaving electrophysiology in dmPFC; n=10). Following arrival at the vivarium, animals were acclimated for 3 days. A 12-h reverse light/dark cycle with lights off at 7:00 AM was utilized. Following acclimation, animals were single housed and given at least a week prior to testing. All animals were at least 70 days of age prior to testing and had *ad lib* access to food and water prior to food restriction/habituation. Animals were food restricted to 85 % of their starting free-feeding weight and maintained under this condition throughout all experiments except immediately prior to and after surgery. All procedures were approved by the IUPUI School of Science Institutional Animal Care and Use Committee and were in accordance with the National Institutes of Health Guidelines for the Care and Use of Laboratory Animals.

### Operant Apparatuses

Eight standard one-compartment operant boxes (20.3cm × 15.9cm × 21.3cm; Med Associates, St Albans, VT) inside of sound attenuating chambers (ENV-018M; MED Associates, St. Albans, VT) were used for both Experiment 1 & 2 Habituation and Shaping protocols. Each box contained left and right retractable levers on one wall, left and right stimulus lights positioned immediately above each lever, and an easily accessible pellet hopper positioned between these left and right positioned devices. The opposite wall contained a house light and a tone generator (2900 Hz) on the topmost position. One custom-built operant box (21.6cm x 25.7cm x 52.0cm) was used to accommodate all electrophysiological experiments. Dimensions, stimuli (including house and cue lights), and retractable levers were all positioned to replicate the conditions of the standard operant boxes as closely as possible. The floor bars of the custom-built box were made of wood polls rather than metal and all metal components of the box were covered in a powder coating to reduce artifacts. In addition, two of the eight standard operant chambers were modified for optogenetic Experiment 1 (see Recording and Stimulation Equipment below for additional information).

### Behavioral Procedures

Following single housing, animals were handled daily for a week. Animals were then habituated to the operant chambers in the same manner described in previous work from our group (see Linsenbardt et al., 2016). Following habituation, animals then began the DD procedure.

The within-session adjusting amount DD procedure was a modified version of the procedure performed by Linsenbardt et al. (2017), which was adapted from Oberlin & Grahame (2009) and is illustrated in (see Figure 1A). Stimuli presentation for a given trial is depicted in Figure 1A, for additional detail (see Linsenbardt et al., 2016). Before beginning the DD task, the *immediate* and *delay* lever were assigned for each animal. The “delay lever” was assigned to each animal as their non-preferred side. Side preference was determined during shaping. Choosing the delay lever always resulted in the delivery of 6 45mg sucrose pellets following a delay (0, 1, 2, 4, 8, or 16-sec). The “immediate lever” was the opposite lever and its value was variable. It began each session with a value of 3 pellets and would increase following delayed choices to a maximum of 6 or decrease following immediate choices to a minimum of 0.

“Forced trials” were implemented for the immediate and delay levers, where two consecutive responses on the same lever would result in a forced trial for the non-chosen lever on the next trial (e.g., trial 1=immediate choice, trial 2=immediate choice, trial 3=delay forced). Implementation of Forced trials is depicted in Figure 1B. If an animal did not lever press for the forced trial, the forced trial would be presented again on subsequent trials until the lever was pressed. The animal had to eventually make a response on the forced trial in order to return to choice trials. There was no effect of forced trials on the value of the immediate lever.

The session terminated either after 30 choice trials or 35 minutes for Experiment 1 (all delays) and for the 0, 1, and 2-sec delays for Experiment 2 (in the standard operant chambers). When Experiment 2 animals were moved from the standard operant boxes to the custom operant box for the awake-behaving recordings (4 and 8-sec delays), sessions terminated after either 40 choice trials or 45min using 20mg sucrose pellets in order to maximize number of trials obtained while recording. The delays were completed in ascending order (0, 1, 2, 4, 8, 16-sec) with a day off in between the start of each new delay. Eight to twelve sessions were given at the 0-sec delay, four sessions at the 1 and 2-sec delay. For Experiment 1, nine sessions were completed at the 4, 8, and 16-sec delays (see supplementary Table 1) to account for optogenetic manipulation. For Experiment 2, recordings during the 4 and 8-s delay were obtained until a viable signal was no longer apparent. Magnitude discrimination was determined at the 0-sec delay in the standard operant chambers using the 45mg sucrose pellets before animals were accepted for surgery with an exclusion criterion of 80% (4.8 pellets) of the maximum reward value (6 pellets) for Experiment 1 and 70% criterion for Experiment 2 (4.2 pellets). The average value of the immediate lever over the last ten choice trials was determined for the last three days of the 0, 1, and 2-sec delay and was used to determine the indifference point of each animal. Animals that were included then either received surgery for Experiment 1 or Experiment 2 (see Surgical Preparation & Implantation for detail).

Indifference points for both Experiment 1 and 2 were obtained by taking the last 10 choice trials of each session for a given delay. For Experiment 1 during ‘experimental delays’ with optogenetic manipulation (i.e., the 4, 8, and 16-sec delays) the last 10 trials of each day for each condition (No inactivation, Epoch 1 inactivation, and Epoch 2 inactivation) were used to determine an indifference point for each condition at each experimental delay (see supplementary Table 1). Days 1 and 2 were excluded for the No inactivation condition, as animals were becoming familiar with the new delay and indifferences points were not yet stable. Therefore, the last 10 trials of days 4, 6, and 8 were used for calculating indifference points for the No inactivation condition. The last 10 trials of days where Epoch 1 as well as Epoch 2 inactivation occurred were taken for each animal for indifference points on optogenetic manipulation days (see supplementary table 1 for experimental design). The rate of discounting was determined using the Mazur Hyperbolic model (equation 2)^55^;

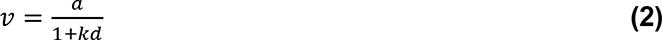

Here, *v* represents the subjective value of the reward, *a* is the fixed value of the delay reward (6 pellets), *d* is the length of the delay (4, 8, or 16-sec), and *k* is the value fitted to the hyperbolic function using Least squares regression to the indifference points at the 4, 8, and 16-sec delays.

These delays were the focus of this analysis as no optogenetic manipulation occurred at the 1, or 2-sec delay. The virus was allowed to express for at least three weeks before beginning any optogenetic manipulation.

### Surgical Preparation & Implantation

For all surgeries, animals were placed inside a flow box and anaesthetized with isoflurane gas (2% at 4L/h) until sedated, at which point they were placed in a stereotaxic frame and maintained on 0.3-0.5% isoflurane for the duration of the surgery. Artificial tears were then applied. Subsequently, fur was shaved and the skin at the incision site was sanitized with three rounds of both 70% EtOH and betadine before applying a local anesthetic (Marcaine; 5mg/kg s.c.). An anti-inflammatory (Ketofen; 5mg/kg dose s.c.) and antibiotic (Cefazolin; 30mg/kg s.c.) were injected at the nape of the neck (anti-inflammatory and antibiotic) before beginning the incision. Once the skull was exposed and cleaned of blood, bregma-lambda coordinates were identified. Prior to any implantation (probe or optic fiber), four stainless steel anchoring screws were inserted. Following insertion of either Cambridge Probes or optic fibers, a two-compound dental cement was used to adhere implants to anchoring screws. Following completion of surgical procedures for Experiment 1 and 2, animals were maintained in a clean heated cage before being returned to the vivarium.

#### Opsin Virus Delivery and Implantation of Optic Fibers (Experiment 1)

Two syringe pumps (Pump 11 Elite; Harvard Apparatus, Holliston, MA) were attached to each arm of the stereotaxic frame and loaded with 2μL Hamilton syringes (7002KH, Hamilton Co., Reno, NV). Coordinates for PL dmPFC viral injections occurred at a 20- degree angle and were as follows: +3.2mm AP, +2.0mm ML, −5.2mm DV from Bregma. Holes were drilled into the skull to allow the Hamilton syringes to penetrate the brain tissue. Animals then received bilateral injections of .65μL at a flow rate of .2μL/min of the inhibitory Adeno-associated virus (AAV-CaMKIIa-eArchT3.0-EYFP; K. Deisseroth via UNC Vector Core) followed by 10 minutes of diffusion before retracting the Hamilton syringes. Subsequently, animals received fiber implantation of Dual Fiber-optic cannulas with guiding sockets (DFC_200/245- 0.37_3.3mm_GS1.4_FLT; Doric Lenses Inc., Quebec, QC, Canada).

#### Electrophysiological Probe Implantation (Experiment 2)

A rectangular craniotomy was performed over the right hemisphere of dmPFC (AP: 2.8, ML: 0.3 from bregma) followed by a durotomy and cleaning/hydration of the probe insertion site with a sterile saline solution. Additionally, two ground screws were placed above the cerebellum. A Cambridge Neurotech F (n=5), P (n=4), or E-series (n=1) 64-channel silicon probe on a movable drive (Cambridge Neurotech, Cambridge, UK) was lowered to the target site. Mobility of the movable drive was maintained with a coating of antibiotic ointment.

### Stimulation and Recording Equipment

#### Optogenetic Stimulation

A green (532nm) laser (MGL-FN-532-300mW; Ultralasers Inc., Toronto, Canada) operated through Med Associates Programming via a TTL (Med Associates, St Albans, VT) was utilized for stimulation. From the fiber coupler, a mono patch cord (MFP_200/240/900- 0.22_1m_FC-FC; Doric Lenses Inc., Quebec, QC, Canada) was attached and traversed the sound attenuating chambers terminating at the rotary joint (FRJ_1×1_FC-FC; Doric Lenses) which attached a Branching Fiberoptic Patchcord (BFP(2)_200/240/ARMO-0.22_0.5m_FCM-GS1.4; Doric Lenses) that was the terminal connection to the animal via guiding socket at the top of the animal’s skull. Stimulation did not occur in pulses and remained on for the duration of the epoch to prevent rebound depolarization of cells. Stimulation at the tip of the fiber measured approximately 21mW resulting in predicted irradiance of ∼60mW/mm2 at the fiber tip. Larger irradiance values were opted for in order to traverse the entire PL cortex with only one fiber per hemisphere.

Stimulation occurred at one of two different epochs during the task for a given session (Epoch 1 inactivation or Epoch 2 inactivation). Epoch 1 stimulation occurred from the start of a given trial and terminated once an animal initiated the trial (see Figure 1A, top). Stimulation remained on if the animal omitted initiating the trial until a response on an initiation lever was made. Epoch 2 stimulation occurred as soon as the animal initiated a trial and terminated once a choice was made (see Figure 1A, top). Stimulation remained if the choice was omitted until a choice was made on subsequent trials. Stimulation occurred on the third, fifth, seventh, and ninth session/day of the 4, 8, and 16-second delays in order to control for carry over effects of the stimulation as well as to obtain indifference points for the No Inactivation condition. All animals received stimulation at both Epoch 1 and Epoch 2 in a cross-over design (see supplementary table 1) so that half the animals received Epoch 1 on the third and seventh day and Epoch 2 on the fifth and ninth day and the other half of animals received the opposite configuration.

#### Electrophysiology Equipment

Silicon probes were acquired from Cambridge Neurotech (Cambridge, UK) and interfaced with Omnetics connectors (Omnetics – Minnesota, US). Silicon electrodes were mounted the day prior to surgery to Cambridge Neurotech microdrives. An Intan RHD SPI cable (Intan – CA) connected the headstage to a Doric Commutator (Doric Lenses – Canada) positioned above the operant apparatus. An OpenEphys (OpenEphys – MA) acquisition system was used to collect all electrophysiological data. AnyMaze (ANY-maze Behavioral tracking software – UK) was used to collect all behavioral and locomotor data. ANY-maze locomotor data was synchronized with OpenEphys via an ANY-maze AMI connected to an OpenEphys ADC I/O board. Med PC behavioral events were also synchronized to the electrophysiological recordings via an OpenEphys ADC I/O board. Following sessions with diminished signal, electrodes were lowered 50µm following completion of that session in order to allow any drifting of the probe to occur before the next day’s session.

### Immunohistochemistry & Histology

#### Optogenetics (Experiment 1)

Animals were perfused within 14 days after behavioral testing with 4% PFA after receiving a anesthetic dose of urethane (1.5-2.0g/kg). Brains were then fixed in 4% PFA for 24 hours before being placed in a 30% sucrose solution (24-72 hours) and subsequently stored at −20 degrees Celsius until sliced 50 microns thick. In order to assess transduction of glutamatergic pyramidal cells within dmPFC, slices were mounted on gelatin subbed glass slides using an aqueous mounting medium (H-1000-10; Vectashield, Invitrogen). A florescence imaging scope (Nikon Eclipse 80*i*; Melville, NY) was used to verify EYFP-tagged protein expression.

#### Electrophysiology (Experiment 2)

Animals were anesthetized with urethane (1.5 – 2.0 g/kg) and subsequently perfused following with 4% PFA after cessation of spinal reflexes. Following tissue extraction, brains were fixed in 4% PFA for 24 hours and then transferred to a 30% sucrose solution for cryoprotection. Following our post-fix procedures, tissue was stored at −80 degrees Celsius until Experiment 2’s cohort was complete. Tissue was sliced at 50 microns and stained for both GFAP and DAPI. Briefly, tissue sections were washed in phosphate buffered saline (PBS) once. Following this, sections were washed in PBS and 0.1% Triton 100. Sections were blocked in 1% normal goat serum. Following blocking, the primary antibody (GFAP; goat anti-chicken) was added and allowed to incubate while shaking for 24 hours at 4°C. Tissue was washed 3x in PBS and then the secondary antibody was added (Alexa fluor 555; goat anti-chicken). Tissue was incubated and shook in a light-protected box for 2 hours at room temperature. Tissue sections were subsequently handled under light-protective materials. Three additional washes in PBS were followed by the addition of DAPI which was allowed to incubate for 10 minutes at room temperature. Three additional washes in PBS followed. Sections were then mounted on gelatin subbed glass slides with anti-fade mounting medium (sc-516212 Santa Cruz Biotechnology) and imaged in order to confirm placement across the dmPFC. Sections were mounted on gelatin subbed glass slides and then imaged in order to confirm placement across the dmPFC.

### Spike sorting

Putative neurons were organized into clusters by Kilosort 2^62^. Following automatic spike-sorting, we determined which of these putative neurons met a qualitative criterion in Phy2 (https://github.com/cortex-lab/phy). Specifically, we ensured that the autocorrelograms contained no refractory violations, we ensured that our waveforms were characteristic of an action potential, and that our signal was minimally contaminated by any noise artifacts. Following qualitative characterization in Phy2, we imported our data into MATLAB (Nantucket, MA) for subsequent analyses. A custom MATLAB routine was used to align spike trains to task events. Spike trains were smoothed using Gaussian convolution with a bin width of 200ms and σ set to 10ms.

### Data Analysis and Statistical Procedures

Data were structured using custom MATLAB routines and subsequently analyzed and graphed using MATLAB for both Experiment 1 and 2. Analysis of initiation and choice latencies used medians, as medians are less sensitive to means to the positive skew of reaction times.

Decision classes (class 1-4) were constructed to parse decisions into four categories based off choice (immediate or delay) and i-value (high or low; see Figure 4B, supplementary Figure 1). Distribution of decision classes were analyzed using a probability density function (PDF) for Laser ON vs OFF conditions (Experiment 1; Figure 4B) and electrophysiological animals (Experiment 2; supplementary Figure 1) to determine how consecutive choices are made within each class. Class 1 and 4 were considered disadvantageous and therefore animals should deviate from exploiting the immediate (class 1) or delay lever (class 4).

To better understand how animals deviate from exploitation strategies, three or more consecutive choices on either the immediate or delay lever were defined as either *immediate exploitation* or a *delay exploitation* strategy, respectively (Figure 1B). Three consecutive choices were defined as exploitation given that an animal continued to choose the same lever despite being exposed to the other lever via a forced trial. Determining whether differences in neural activity exist for when animals continued to exploit a choice option (e.g., four choices on the same lever) vs deviate from exploitation (e.g., three consecutive choices followed by a choice on the opposite lever) were then analyzed using PCA. Continued exploitation of a choice option was defined as either Immediate-Fail-to-Change or Delay-Fail-to-Change whereas deviating from the exploit strategy was defined as either Immediate-Change or Delay-Change, resulting in four strategy conditions to be analyzed (see Figures 6-8).

PCA was run separately for the 4-sec and 8-sec delays. For all PCA analyses, spike trains were aligned to the choice point comprised of an interval of 19 seconds prior to and 1 second after the choice (−19s to +1s). Using the choice-point-aligned spike trains, for each neuron, the firing rates (FR) were normalized by the mean firing rate of that unit. Each PCA analysis consisted of 101 time bins (200ms each bin). To be included in the PCA analysis, only neurons firing in all eight trial types (see Figure 6A key for trial types) that occur during trials three and four of the four possible strategy conditions were included in the analysis. Firing rates were normalized and smoothed using a moving average filter spanning 5 bins for each of the eight trial types (trials three and four for each of the four strategy conditions). The eight vectors corresponding to the eight trial types were then concatenated and z-scored prior to running PCA. FR from each neuron (n=581 4-sec, n=1166 8-sec) were contained in columns and each of the 101 time bins per trial type in rows. For example, the for the 4-sec delay, the matrix *F*(808 × 581) contained n=581 neurons and each column and rows consisted of eight blocks containing the 101 timepoints for each condition (total of 808). PCA was then conducted to analyzed neural activity across all 101 time bins for trials three and four of each strategy condition. The 3 most explanatory dimensions were chosen (top 3 PCs).

PCA for each individual session (4-sec or 8-sec delay) was conducted to obtain trajectories for each of the eight trial types. Using the trajectory coordinates derived from individual session PCA, the mean Euclidean distance in PCA state-space was calculated between the 3^rd^ and 4^th^ trial of the series for each strategy condition. PCA for the choice classes (1-4) were conducted in the same manner. The only factor that differed in this analysis was that only four trial types were present for each PCA conducted, given that each class only contained one of the possible two choice alternatives (Immediate or Delay). Therefore, class 1-2 PCA contained four trial types (3^rd^ and 4^th^ trials) corresponded to either the Immediate Change or Fail-to-Change strategies, while class 3-4 contained the 3^rd^ and 4^th^ trial corresponding to the Delay Change or Fail-to-Change strategy conditions. The number of neurons used within each PCA analysis is depicted in Supplementary Table 2.

**Table S1:** **Schedule of optogenetic inhibition for each experimental delay (4, 8, 16-sec) for Experiment 1.** Each **X** indicates a Laser OFF day. The 1 and 2 indicate Laser ON days (Epoch 1 or 2 inactivation; see Figure 1A). Epochs 1 and 2 were alternated every other day such that each animal received two ‘Laser ON days’ at both Epoch 1 and 2. See Figure 1 for Epoch 1 and 2 duration prior to choice. Days 1 and 2 were discarded from analyses as animals were adjusting to the new delay and behavior was unstable.

Tables/Figures
|  | Day 1 | Day 2 | Day 3 | Day 4 | Day 5 | Day 6 | Day 7 | Day 8 | Day 9 |
| --- | --- | --- | --- | --- | --- | --- | --- | --- | --- |
| A | X | X | 1 | X | 2 | X | 1 | X | 2 |
| B | X | X | 2 | X | 1 | X | 2 | X | 1 |

**Table S2.** Number of neurons and session numbers used in each PCA analysis. The total neurons, excluded neurons, and end number of neurons are included. Neurons were excluded for not firing in all trial conditions tested in the PCA analysis. See Materials and Methods for exclusion criteria.

| Delay | Starting Neuron Number | Number Excluded Neurons | Total neurons used for PCA | Session Numbers Used for PCA Analysis |
| --- | --- | --- | --- | --- |
| 4s | 2120 | 1539 | 581 | 1, 4, 7, 9, 15, 16, 21, 44, 45, 48, 50 |
| 8s | 2078 | 912 | 1166 | 23, 28, 29, 30, 31, 32, 43, 52, 53, 54, 55, 57, 58 |

**Figure S1.**
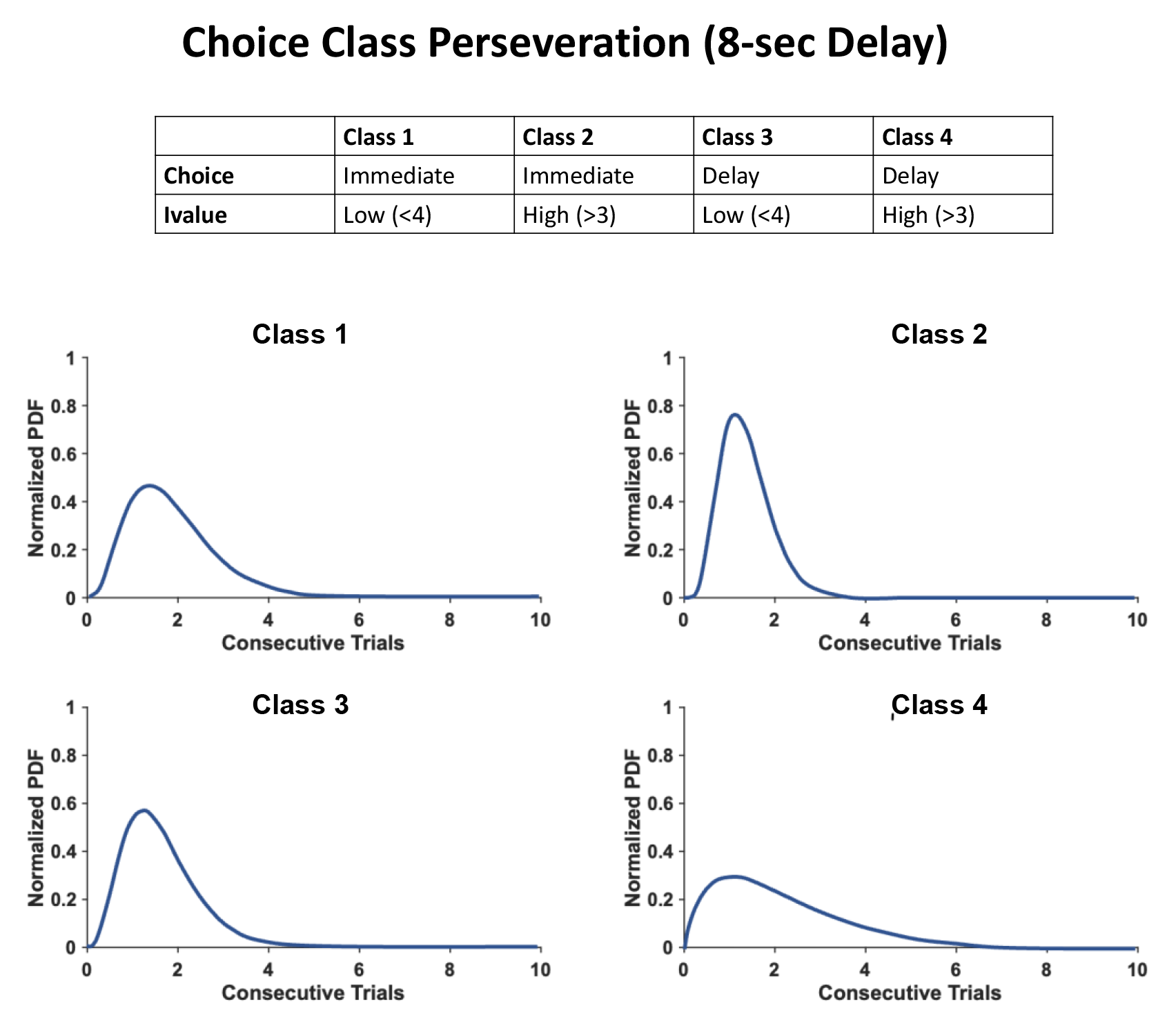
Experiment 2 awake-behaving electrophysiology animals normalized probability density function distribution of consecutive choices for each individual choice class.

**Figure S2.**
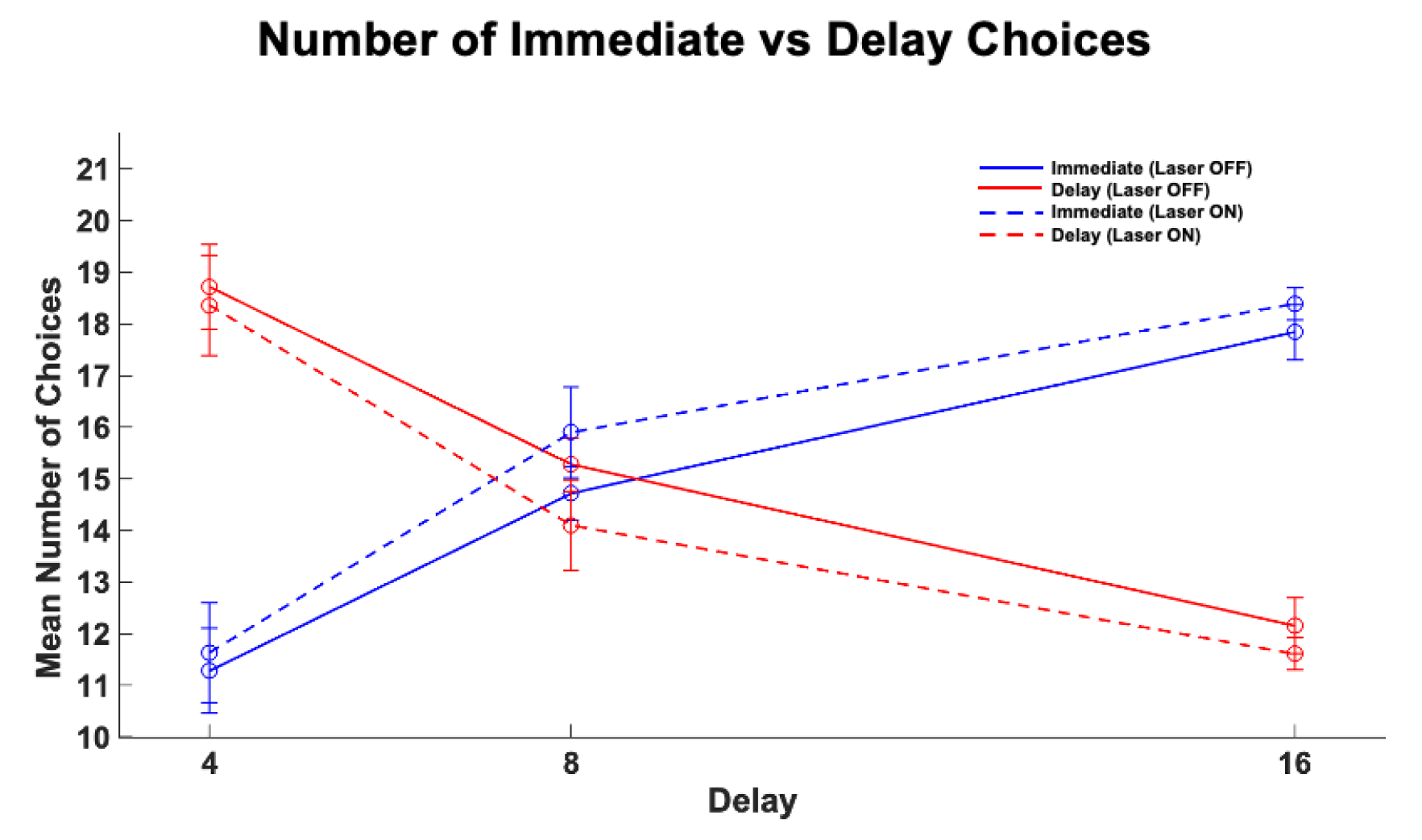
Number of Immediate and Delay choices made during the 4, 8, and 16-sec delays for Experiment 1 optogenetic animals during Laser ON and Laser OFF sessions. Sessions where no optogenetic manipulation (Laser OFF) sessions are depicted using dashed lines while optogenetic manipulation sessions (Laser ON) are depicted by solid lines. Mean and +/- SEM are depicted for each delay. Statistically significant interaction between Choice Lever (IM or DEL) and Delay (2-way ANOVA: F(2,191) = 52.43, *p* < 0.001). For Laser OFF sessions, the number of Immediate and Delay choices did not differ at the 8-sec delay (p >.05), however the number of Delay choices were greater at the 4-sec delay (p < .001) and Immediate choices were more numerous at the 16-sec delay (p < .001), as indicated by a Scheffe multiple comparison test.

**Figure S3.**
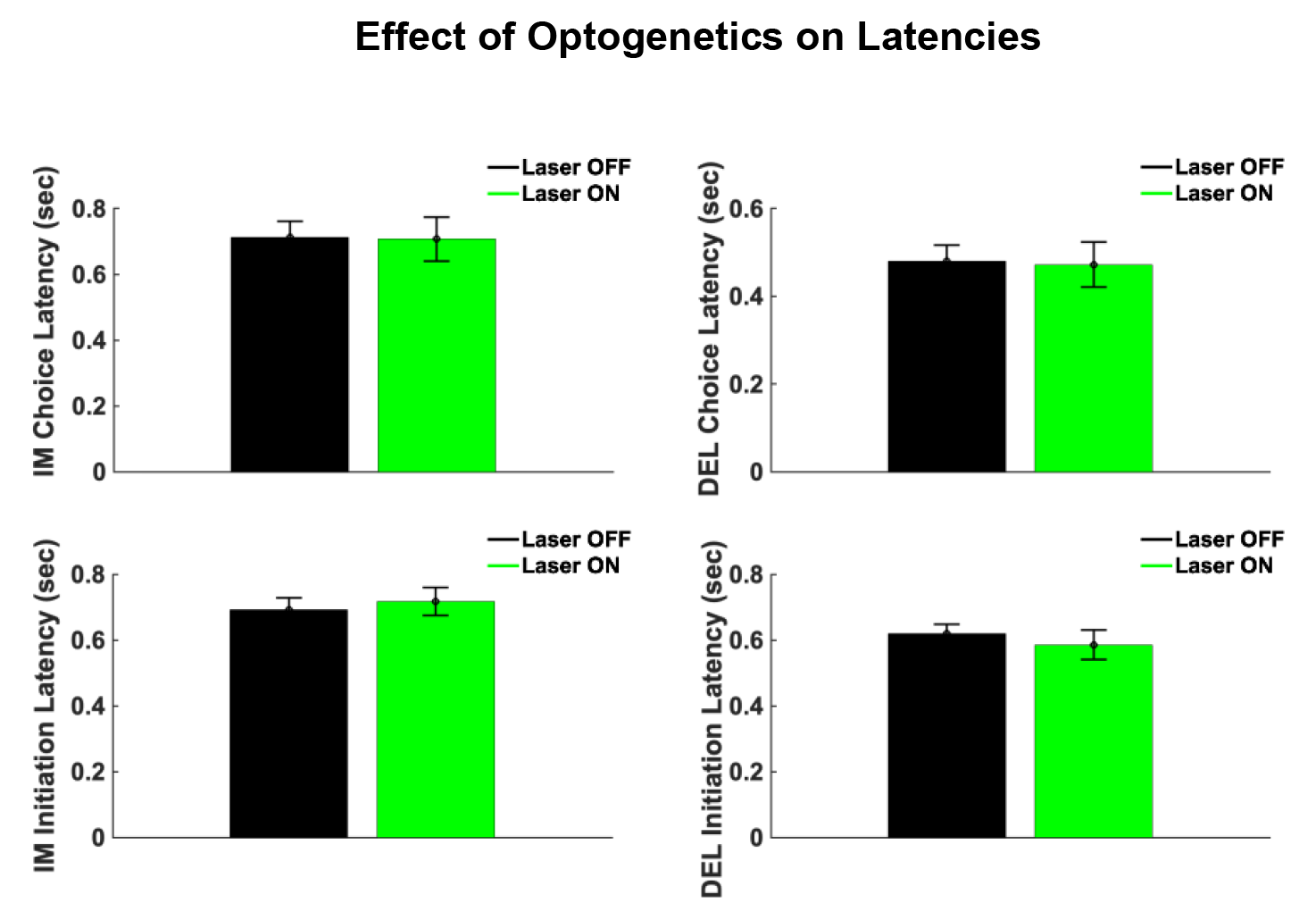
No effect of optogenetic inhibition of dmPFC on initiation or choice latencies (Median +/- SEM) for either Immediate (IM) or Delay (DEL) choices (Wilcoxon Rank Sum test: *Z*<.70, *p*>.48).

